# Multitask Knowledge-primed Neural Network for Predicting Missing Metadata and Host Phenotype based on Human Microbiome

**DOI:** 10.1101/2024.02.05.578930

**Authors:** Mahsa Monshizadeh, Yuhui Hong, Yuzhen Ye

**Author notes:** These two authors contribute equally to the paper (Mahsa Monshizadeh, Yuhui Hong).

## Abstract

Microbial signatures in the human microbiome have been linked to various human diseases, and Machine Learning (ML) models have been developed for microbiome-based disease prediction, although improvements remain to be made in accuracy, reproducibility and interpretability. On the other hand, confounding factors, including host’s gender, age and BMI can have a significant impact on human’s microbiome, complicating microbiome-based human phenotype predictions. We recently developed MicroKPNN, an interpretable ML model that achieved promising performance for human disease prediction based on microbiome data. MicroKPNN explicitly incorporates prior knowledge of microbial species into the neural network. Here we developed MicroKPNN-MT a unified model for predicting human phenotype based on microbiome data, as well as additional metadata including age, body mass index (BMI), gender and body site. In MicroKPNN-MT, the metadata information, when available, will be used as additional input features for prediction, or otherwise will be predicted from microbiome data using additional decoders in the model. We applied MicroKPNN-MT to microbiome data collected in mBodyMap, covering healthy individuals and 25 different diseases, and demonstrated its potential as a predictive tool for multiple diseases, which at the same time provided predictions for much of the missing metadata (e.g., the BMI information was missing for 94% of the samples). Our results showed that incorporating real or predicted metadata helped improve the accuracy of disease predictions, and more importantly, helped improve the generalizability of the predictive models. Finally, our model enables the interpretation of predictive models and the identification of potential microbial markers affecting host phenotypes.

## Introduction

The human microbiome is a sophisticated ecosystem encompassing trillions of microorganisms, demonstrating a crucial impact on human health and diseases [1]. This intricate microbial community is distributed across various body sites, including the skin, oral cavity, respiratory tract, gastrointestinal tract, urinary tract, and reproductive tract [2]. It is involved in diverse physiological processes, ranging from digestion and immunity to metabolism and brain function. Maintaining the delicate balance of the microbiome is essential, as disruptions can result in dysbiosis, associated with a broad spectrum of diseases, including inflammatory conditions, obesity, diabetes, and depression. Consequently, the exploration of the human microbiome has emerged as a rapidly expanding field of research, holding substantial promise for advancements in disease diagnosis and treatment, among others. Metagenomic data analysis of the human microbiome has become a widely adopted strategy for exploring its impact on human health and disease. The link between the human microbiome and human health has generated considerable interest in using microbiome data for disease diagnosis and personalized medicine [3].

Many different predictive models have been developed for microbiome-based disease prediction [4, 5]. The models vary in the types of inputs they take (species profiles, functional profiles, or both), Machine Learning (ML) and AI algorithms, and the prediction targets (single-disease or multi-disease) [6, 7]. These models have various prediction accuracy and interpretability (some are black box whereas others are more interpretable). Early predictive models were based on conventional ML/AI algorithms, and more recently, deep learning methods including various autoencoders were also exploited for learning the representation of quantitative microbiome profiles in a lower dimensional latent space for building predictive models. Examples include DeepMicro [8], Ph-CNN [9], PopPhy-CNN [10], and EPCNN [11]. We have previously developed MicroKPNN [12] for predicting human health status based on microbiome data, aiming to improve the accuracy of prediction and at the same time provide good explainability of the predictions. MicroKPNN incorporates multiple microbial relationships (metabolic, phylogenetic, and community) in the model to improve the performance of microbiome-based prediction and interpretability of the models. The model enables the examination of the importance of different input species and possible explanations (through the hidden nodes that have biological meaning). MicroKPNN achieved encouraging performance when tested on seven gut microbiome datasets involving five different human diseases.

Predictions of human diseases that could benefit from using microbiome include liver diseases [13], respiratory diseases (asthma [14] and COPD [15]), diabetes (T1D [16] and T2D [17]), inflammatory bowel disease [18], and neuropsychiatric conditions [5]. A growing body of work has also demonstrated the possibility of using the gut microbiome for the prediction of future onset of diseases. Examples include a longitudinal study showing that the gut microbiome at 1 year of age can distinguish individuals who develop future T1D up to 20 years later from those who do not [19], and another study that shows baseline gut microbiome is associated with new-onset T2D in up to 18 years [20].

Various confounding factors affect human microbiota, and the factors include sex, diet, race, medications, host’s genetic variation, and so on. In addition, there are technical artifacts that could complicate the use of microbiome data [21]. Animal and human studies have shown sex differences in gut microbiota, and different mechanisms were suggested [22, 23]. Results from an animal experiment involving microbiota transfer suggested that the microbiota-independent gender differences in the immune system select a gender-specific gut microbiota composition, which in turn further contributes to gender differences in the immune system [24]. Sex hormones are a potent driver of differences in the microbiome, and other factors (diets, antibiotics and environment) impact gut microbiota in a sex-dependent manner [23]. A joint analysis [25] of the composition of the human microbiome and host genetic variation revealed significant associations between host genetic variation and microbiome composition, and these associations are found to be driven by host genetic variation in immunity-related pathways and genes associated with microbiome-related complex diseases including inflammatory bowel disease and obesity-related disorders.

In addition to models that have been developed to utilize microbiome data for human disease prediction, microbiome-based ML models have been developed for other applications, including age prediction [26, 11]. ML models based on random forest regression revealed different levels of accuracy using human skin, oral, and gut microbiomes, with the model using the skin microbiome achieving the most accurate age prediction [26]. A multi-view learning based model was developed for age prediction using compositional and functional features of microbiome [11]. A model was built for predicting body mass index (BMI) using microbiome data [27], achieving BMI predictions with a mean absolute error (MAE) of about 2 *kg/m*^2^. The human microbiome may be used as a resource in the forensics toolkit [28], as human-associated bacterial DNA can be used to uniquely identify an individual [29], and even to provide information about their life and behavioral patterns [30].

Considering that microbiome composition reflects the impacts of many factors on the microbiota, and is associated with host phenotypes, here we propose a unified predictive model for the various metadata and human diseases. Although our focus is on microbiome-based disease prediction, our model also predicts the metadata including age, gender, BMI, and body site if they are missing. We showed that by doing this, we can not only improve the accuracy of disease prediction, but also improve the generalizability of the predictive model. We called our new model MicroKPNN-MT as it incorporates prior-knowledge as in MicroKPNN [12]. In addition, MicroKPNN-MT is a multitask and multiclass classification model: it includes individual decoders for metadata predictions, and the decoder for disease prediction is multiclass. Since most metagenomics projects studying microbiome-disease association often have healthy samples (control), our multiclass classification model enables the utilization of all these healthy samples from different projects. We applied our model to the mBodyMap dataset [31], covering 25 different human diseases, and demonstrated its potential as a predictive tool for multiple diseases and identifying microbial markers for the diseases and other metadata such as age.

## Materials and Methods

### Human microbiome data

We used the mBodyMap database [31], which offers a comprehensive collection of human metagenomic data and their species abundance profiles derived using state-of-the-art tools. Reads processing and taxonomic assignments for all the datasets included in mBodyMap were done using the same set of tools [31]. For taxonomic assignments, MAPseq (v1.2) [32] was used for 16S rRNA sequencing data, and MetaPhlAn2 [33] was used for shotgun metagenomic sequences. The database also boasts a carefully curated set of human-related metadata, including information on diseases and health.

We excluded the samples with the sum of relative abundances *<* 90 (%), and excluded diseases that had *<* 50 samples. We ended up with 34,233 samples from 56 projects, involving 25 diseases. A total of 6,052 species were identified from these samples. All samples have body site information, whereas other metadata have various levels of incompleteness: 6,422 samples have age information, 23,804 samples have gender information, and only 2,094 samples have BMI details.

Our experiments showed that categorized age and BMI worked better for our application compared to their actual values. So, we performed preprocessing on the collected data. Specifically, we categorized BMI data qualitatively based on standard definitions: underweight (BMI *<* 18.5), healthy weight (18.5 *≤* BMI *<* 25), overweight (25 *≤* BMI *<* 30), and obesity (BMI *≥* 30). Additionally, age data was grouped into qualitative categories including infant (age *≤* 3), children adolescents (3 *<* age *≤* 18), young adult (18 *<* age *≤* 35), middle aged (35 *<* age *≤* 50), senior (50 *<* age *≤* 65), and elderly (age *>* 65). See Figure 1 for the breakdown of the samples according to the different metadata.

**Figure 1:**
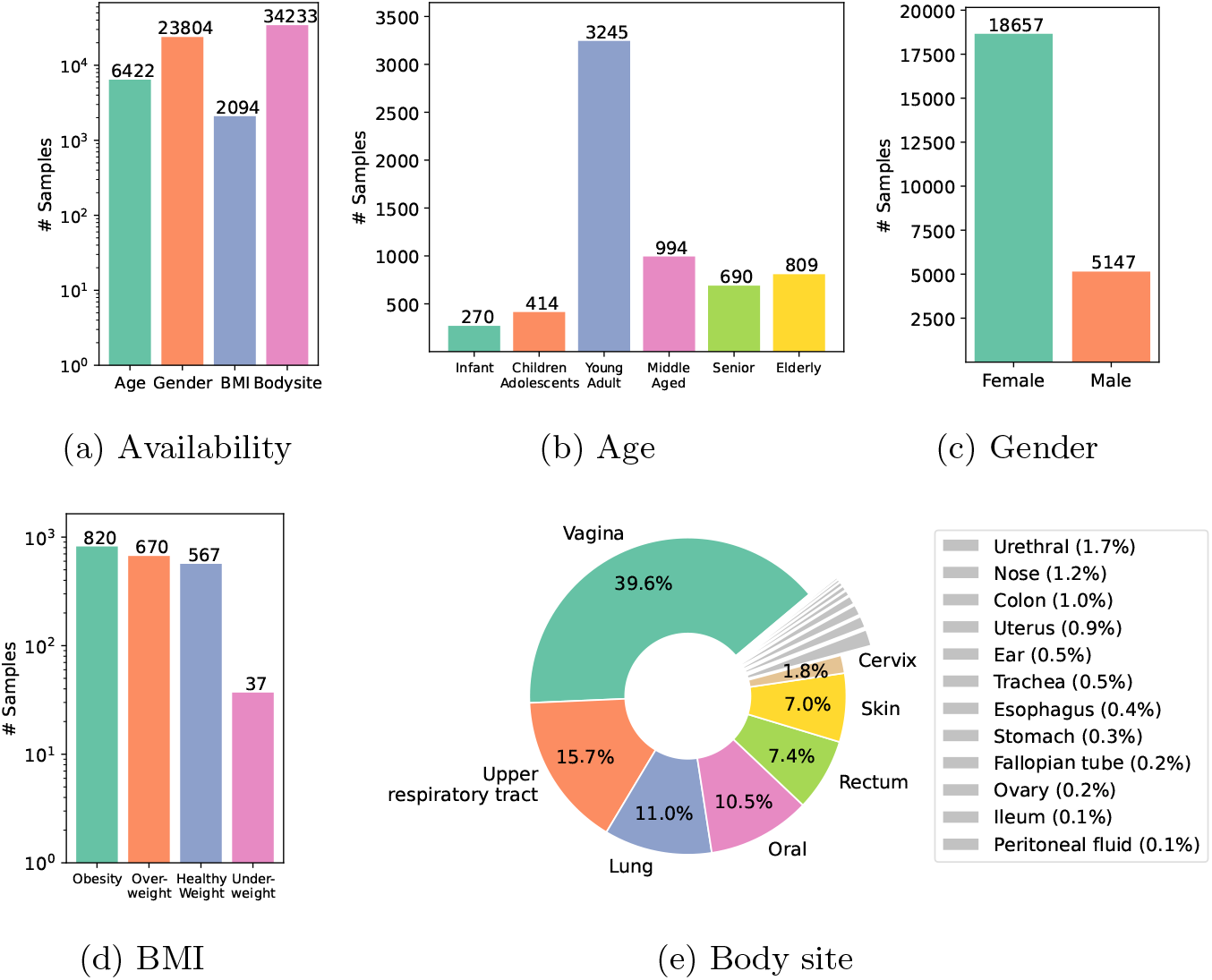
Breakdown of the samples according to the metadata.

### The model architecture

In this study, we propose a novel neural network architecture, termed MicroKPNN-MT, designed to capture complex relationships within human microbiome samples. The architecture is tailored to incorporate prior knowledge about microbial interactions, taxonomic relationships, and community structures for microbiome-based human disease prediction. Furthermore, MicroKPNN-MT utilizes metadata (age, gender, BMI, and body site) to enhance disease prediction. For the samples with missing metadata, MicroKPNN-MT predicts the missing metadata using additional decoders in the model, and predicted metadata is used for disease prediction. Figure 2 shows the overall model architecture, which comprises several key components outlined below.

**Figure 2:**
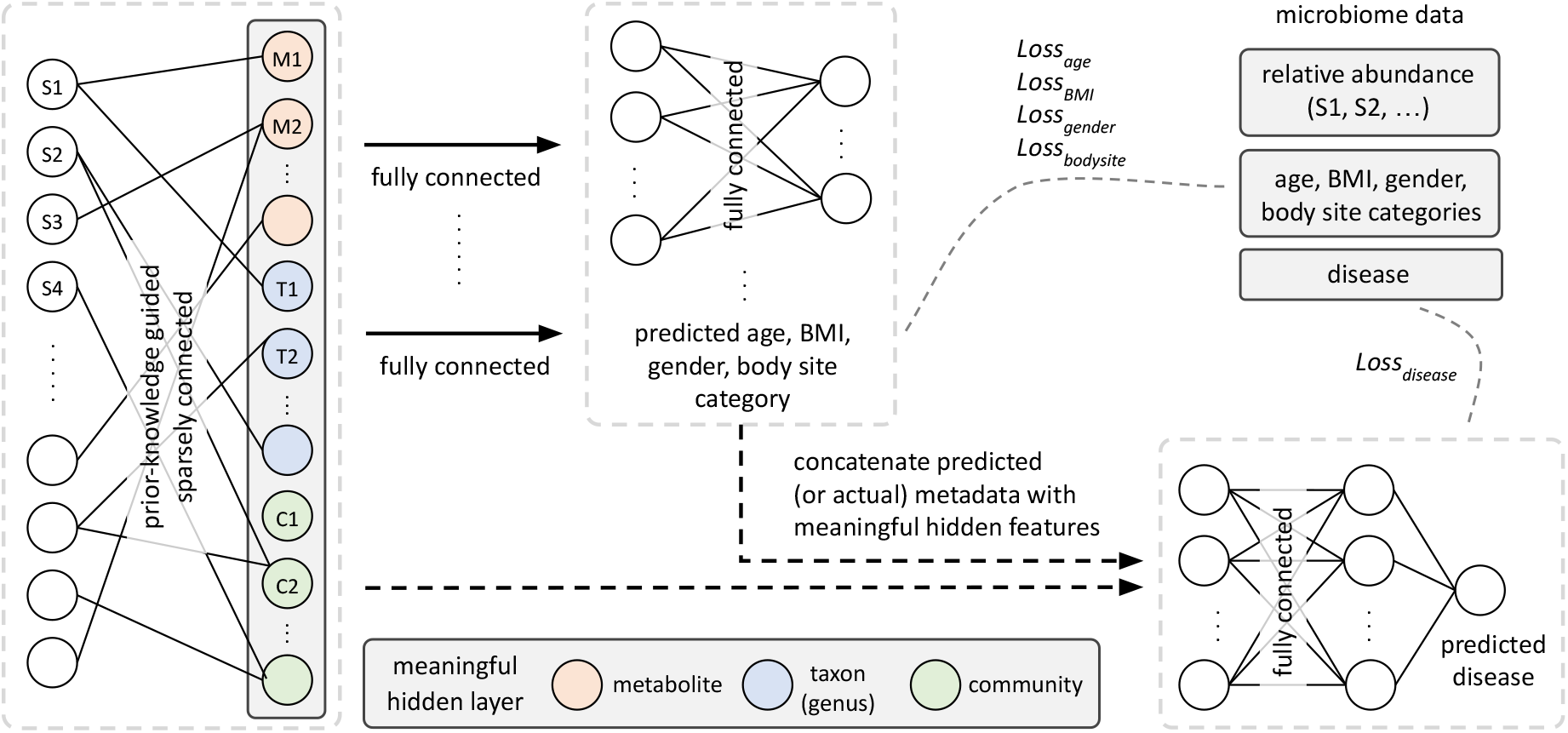
Model architecture used in MicroKPNN-MT for multitask and multiclass classification. It predicts human phenotype and missing metadata based on microbiome data.

#### Input layer

The input layer of the neural network is composed of species abundance data derived from human microbiome samples. Each input node represents the relative abundance of a specific microbial species in the sample. We used the species abundance data provided by mBodyMap [31].

#### Customized linear layer (MaskedLinear)

To leverage prior knowledge effectively, the first hidden layer employs a customized linear layer, denoted as MaskedLinear. This layer enforces a mask on the connections between the input and hidden layers based on prior knowledge. The mask is designed to capture relationships among three distinct groups: metabolites, taxa, and communities.

- Metabolites (pink nodes in Figure 2). Each metabolite is represented by two nodes in the hidden layer: one for production and one for consumption. Edges connect producer species in the input layer to the production node and consumer species to the consumption node. The metabolic edges are created according to the NJS16 metabolic network [34].
- Taxa (blue nodes). Taxonomic relationships are encoded using the NCBI hierarchical taxonomy. Edges connect species in the input layer with their corresponding genus. We note that MicroKPNN could exploit different taxonomic ranks in this hidden layer (genus, order, etc). MicroKPNN-MT uses the genus as the taxonomic rank, as it significantly reduces the time complexity by eliminating the search for taxonomic ranks, and also in MicroKPNN, we showed that using the genus generally gave good results across different datasets and diseases.
- Communities (green nodes). Each node represents a community, and all species belonging to the community have an edge connecting to the node. Communities are inferred from a species co-occurrence network [35].

#### Metadata decoders

A dedicated decoder is included for each metadata (age, gender, BMI, and body site). Each decoder comprises one fully connected layer with dimensions matching the MaskedLinear layer. The output dimensions of these decoders align with the number of classes in each metadata, except for gender, which has a single output with a sigmoid activation function.

#### Disease decoder

An additional decoder is introduced for disease prediction, taking input from the output of the metadata decoders and the customized layer. This decoder includes one fully connected layer with dimensions matching the MaskedLinear layer. The output dimension matches the number of classes in the disease prediction task. To enhance disease prediction accuracy, metadata are masked in the model. If true values for metadata are available, they are used directly. Otherwise predicted metadata from the metadata encoders are utilized for disease prediction.

### Model training and performance evaluation

We adopted a 5-fold cross-validation strategy to evaluate the performance of our model. Special care was taken to ensure that each fold maintained a consistent distribution of diseases, enhancing the reliability of our results.

To balance the effects of metadata that have different numbers of available samples, we use weighted cross-entropy and binary cross-entropy as the loss function:

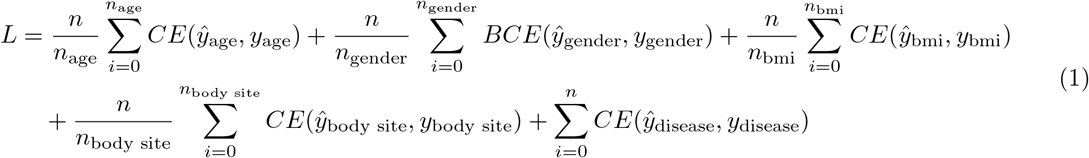

where *n, n*_*age*_, *n*_*gender*_, etc., denote the sample counts for the entire dataset, age, gender, etc., respectively; *y* is the actual category, *ŷ* is predicted category; *CE* and *BCE* represent cross-entropy and binary cross-entropy losses, respectively.

The performance of the model was assessed using the following metrics: accuracy (ACC), which is the fraction of correct predictions; area under the curve (AUC), an aggregated measure of the model’s performance across various decision thresholds; and F1 score, which is the harmonic mean of the precision (fraction of true positives among all predicted positives) and recall scores (fraction of predicted true positives among all true positives). As our model is multiclass involving 25 diseases and healthy individuals for phenotype prediction, the metrics were computed for each phenotype, then averaged to provide the overall scores. The same approach was used for the evaluation of the metadata prediction.

These metrics were chosen to account for the imbalanced nature of the dataset we used, providing a comprehensive assessment of the model’s performance across individual diseases as well as on average.

### Interpretation of the models

We implemented local interpretability methods to analyze individual samples and subsequently aggregated these interpretations into global ones with a particular focus on the metadata-related architecture within MicroKPNN-MT. Specifically, to measure the influence of various metadata elements on disease prediction and the impact of specific hidden nodes on metadata prediction, we employed the Integrated Gradients [36] and Layer Conductance [37, 38], respectively.

Integrated Gradients (IG) attribute NN predictions to input features, while Layer Conductance (LC) delves into individual neuron and layer contributions. IG and LC attributes, positive or negative, reflect a feature’s influence on the prediction. In our experiments, we transform these attributes into importance scores by taking their absolute values, where a higher score indicates a larger impact of a feature on a specific prediction. During the aggregation phase, to determine the overall importance of features to a specific task, we average the scores across all classes within that task. In addition, the model is trained 20 times with randomly initialized weights for a stable and credible interpretation. This approach enables a thorough interpretability analysis of MicroKPNN-MT, exploring both the contributions of the nodes to metadata prediction and the impacts of metadata on disease prediction.

### Implementation and availability

We developed MicroKPNN-MT using PyTorch, and for interpretability analysis, we employed Captum [39], a package specifically designed for model interpretation within the PyTorch framework. We utilized AdamW optimizer with early stopping to mitigate overfitting. The learning rate is set to 0.001 and the batch size is set to 16. The source code of MicroKPNN-MT is available at: https://github.com/mgtools/MicroKPNN-MT.

## Results

### Using metadata helps improve the disease prediction

We conducted a comprehensive comparison between disease prediction models with and without metadata integration. The primary objective was to assess the impact of incorporating metadata on the predictive performance of the models. Despite the consistent structure maintained across all models, the crucial distinction lies in the meaningful interpretation of the metadata nodes within each model.

The results (see Table 1) show a notable improvement in disease prediction outcomes when all metadata were included in the models, with the F1 score improved from 0.763 to 0.814. We also built different models incorporating only one metadata at a time to test the impact of individual metadata on the prediction. The results showed that incorporating actual or predicted metadata, especially age and body site information helped improve disease prediction (see Table 1).

**Table 1:** Comparison of the accuracy of disease prediction with and without using metadata.

| Model | Accuracy (std) | AUC (std) | F1 score (std) |
| --- | --- | --- | --- |
| disease only | 0.913 (0.00357) | 0.966 (0.048) | 0.763 (0.194) |
| disease + age | 0.919 (0.00185) | 0.967 (0.048) | 0.785 (0.180) |
| disease + gender | 0.913 (0.00341) | 0.967 (0.046) | 0.762 (0.196) |
| disease + BMI | 0.911 (0.00093) | 0.965 (0.045) | 0.757 (0.204) |
| disease + body site | 0.920 (0.00441) | 0.968 (0.048) | 0.789 (0.183) |
| disease + all metadata | 0.924 (0.00228) | 0.969 (0.049) | 0.814 (0.169) |

Figure 3 shows that different diseases have a wide range of prediction accuracy. MicroKPNN-MT predicted some diseases including cystic fibrosis with high accuracy. Fewer samples for training probably contributed to the poorer performance of some of the phenotypes including Parkinson disease, Psoriasis, and GPA. On the other hand, it also suggests these diseases may involve more complicated factors, and using microbiome data alone was not sufficient for accurate predictions.

**Figure 3:**
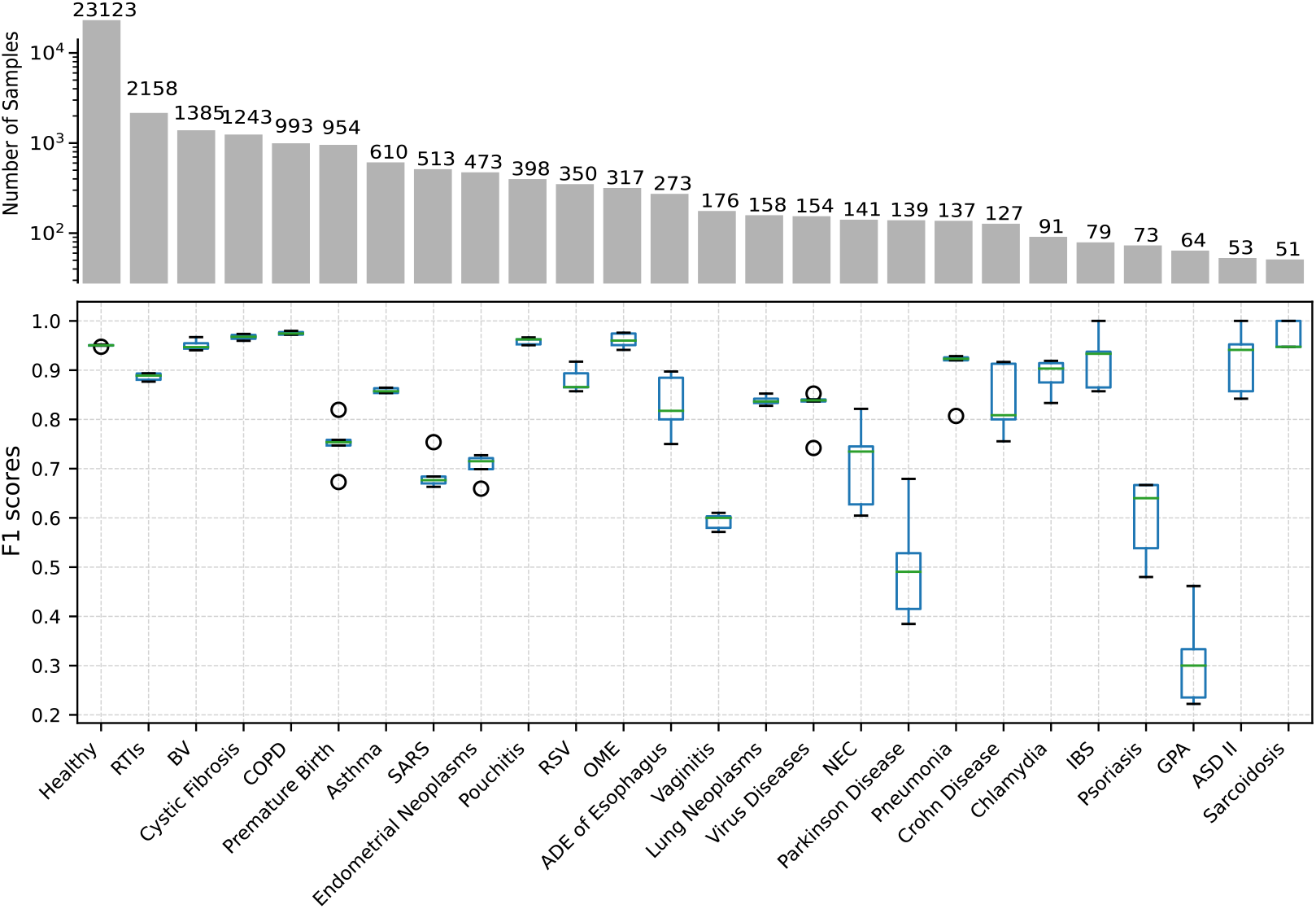
Prediction accuracy for each of the phenotypes (25 diseases + healthy) in F1 scores. The boxes with whiskers show the distribution of the F1 scores (mean and variance computed from the 5-fold cross-validations). The bars on the top show the number of available samples for each phenotype for training and testing. Abbreviations for the diseases: RTIs (Respiratory Tract Infections), BV (Bacterial Vaginosis), COPD (Chronic Obstructive Pulmonary Disease), OME (Otitis Media with Effusion), NEC (Necrotizing Enterocolitis), IBS (Irritable Bowel Syndrome), GPA (Granulomatosis with Polyangiitis).

### Missing metadata can be predicted from microbiome data

As summarized in Table 2, all metadata can be predicted using microbiome data with promising accuracy. For this comparison, for each metadata, we used predictions from two different models, one for disease prediction and all metadata (disease + all metadata), and the other one for disease prediction and the targeted metadata (disease + one metadata). The results show that these two models gave very similar results, although the disease + one metadata model gave marginally better results. We note age has 6 categories (from infant to elderly), BMI has 4, sex has 2, and there are 19 different body sites. BMI prediction has the lowest accuracy of about 0.58 (still significantly better than random guesses involving four choices). We attribute the poor BMI prediction to the small training data (only 6% of the samples have BMI information), among other possible reasons. Figure 4 shows the confusion matrices for the predictions of age and BMI, showing that for most samples, their age and BMI can be correctly predicted, however significant confusion especially between neighboring categories (e.g., elderly and senior, young adult and middle aged, healthy weight and overweight) were observed.

**Table 2:** Summary of the metadata prediction in ACC/AUC/F1 score (std).

| Model | Age |  |  | Gender |  |  | BMI |  |  | Body site |  |  |
| --- | --- | --- | --- | --- | --- | --- | --- | --- | --- | --- | --- | --- |
|  | ACC | AUC | F1 | ACC | AUC | F1 | ACC | AUC | F1 | ACC | AUC | F1 |
| disease + one metadata | 0.765<br>($<0.01$) | 0.886<br>(0.062) | 0.688<br>(0.135) | 0.933<br>($<0.01$) | 0.967<br>($<0.01$) | 0.902<br>(0.055) | 0.591<br>(0.028) | 0.778<br>(0.075) | 0.494<br>(0.187) | 0.937<br>($<0.01$) | 0.944<br>(0.075) | 0.652<br>(0.328) |
| disease + all metadata | 0.759<br>(0.011) | 0.885<br>(0.070) | 0.679<br>(0.140) | 0.932<br>($<0.01$) | 0.967<br>($<0.01$) | 0.901<br>(0.056) | 0.584<br>(0.022) | 0.776<br>(0.072) | 0.467<br>(0.211) | 0.934<br>($<0.01$) | 0.940<br>(0.086) | 0.638<br>(0.339) |

**Figure 4:**
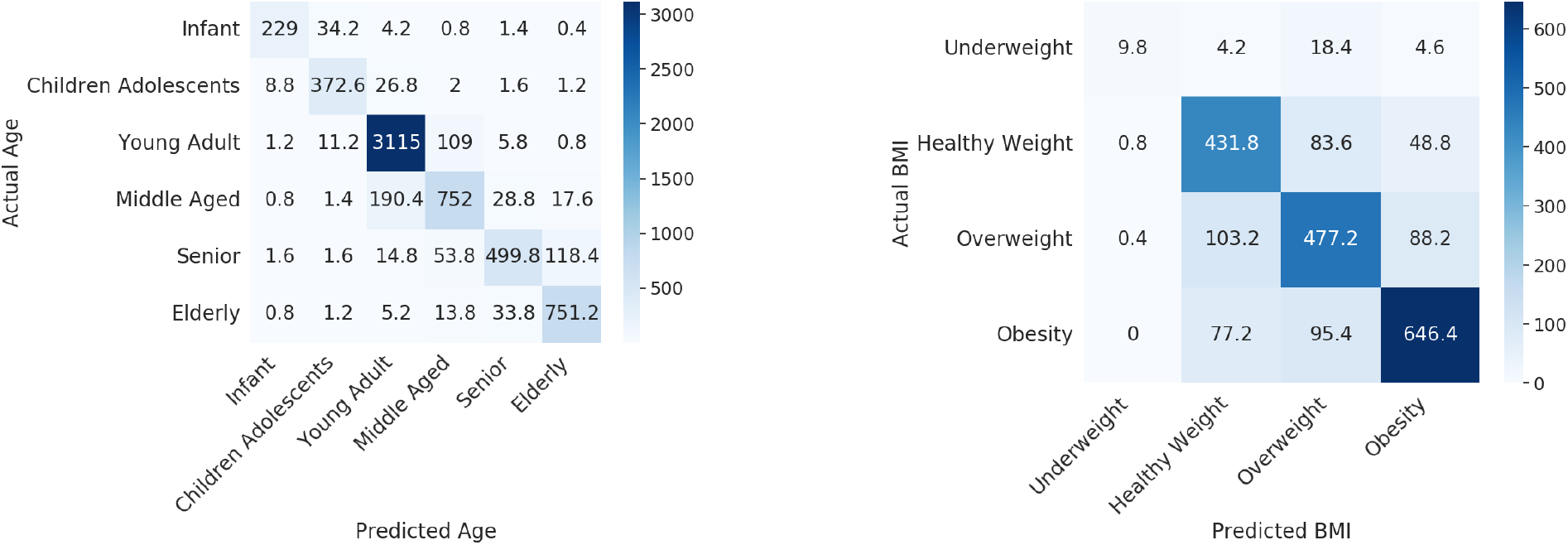
Confusion matrices summarizing the predictions of age and BMI, respectively. The number of samples in each cell was averaged over 5-fold cross-validations.

### Incorporating metadata helps improve the model generalizability

To assess the robustness and generalizability of our model, we designed another experiment so that predictive models were applied to unseen data from projects that were not included in the training. We selected three diseases (out of 25 diseases) that have data from three or more projects. Additionally, we included a set of healthy samples, expanding the disease prediction task to involve four classes, namely Cystic Fibrosis, Chronic Obstructive Pulmonary Disease, Bacterial Vaginosis, and the healthy class.

We divided our dataset into two main subsets: a training dataset and a test dataset. The training dataset comprised samples from 10 projects, totaling 4380 samples. For the test dataset, we selected a different set of 6 projects (no overlaps between projects for train and projects for testing) for a total of 869 samples. This partitioning strategy aimed to provide a comprehensive evaluation of our model’s performance on a diverse range of projects and diseases. We trained and compared two models: one that utilized metadata and another that did not include this additional information.

We examined the performance of the models on the training and test datasets, respectively (see Table 3). On the training dataset, the model using metadata yielded a slightly improved performance, highlighting the potential benefits of incorporating additional contextual information even in familiar data scenarios. By contrast, on the test dataset, the model incorporating metadata consistently outperformed the counterpart that lacked this additional information (F1 score improved from 0.705 to 0.856), showcasing its superior generalizability on previously unseen data. This suggests that the integration of metadata plays a pivotal role in bolstering the model’s predictive capacity, particularly in scenarios where it encounters diverse and unfamiliar data. However, the results suggest there is still room for further improvement for generalization, as we saw a significant performance degradation on the unseen samples, despite that metadata helped.

**Table 3:**
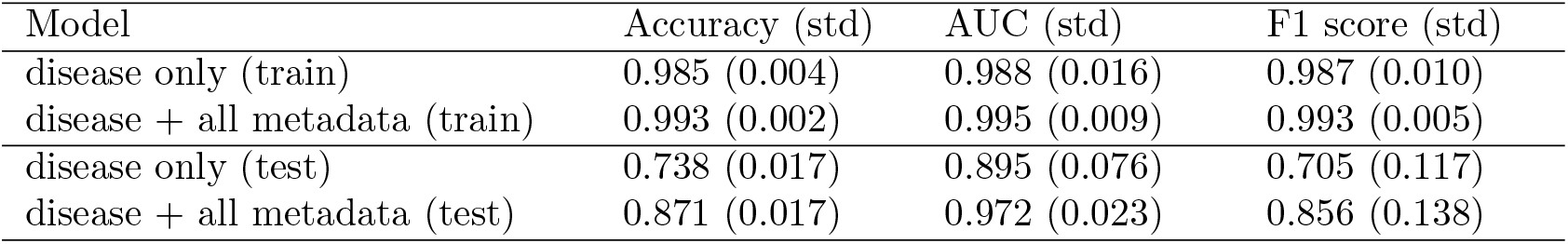
Comparison of the accuracy of disease prediction with and without using metadata on samples from unseen projects (not used for training).

### Interpretation of the predictive models

Importance scores of the metadata and the individual nodes computed from the predictive models can be used to shed light on the impacts of metadata on disease prediction and to explain predictions. Figure 5 shows some example applications of importance scores.

**Figure 5:**
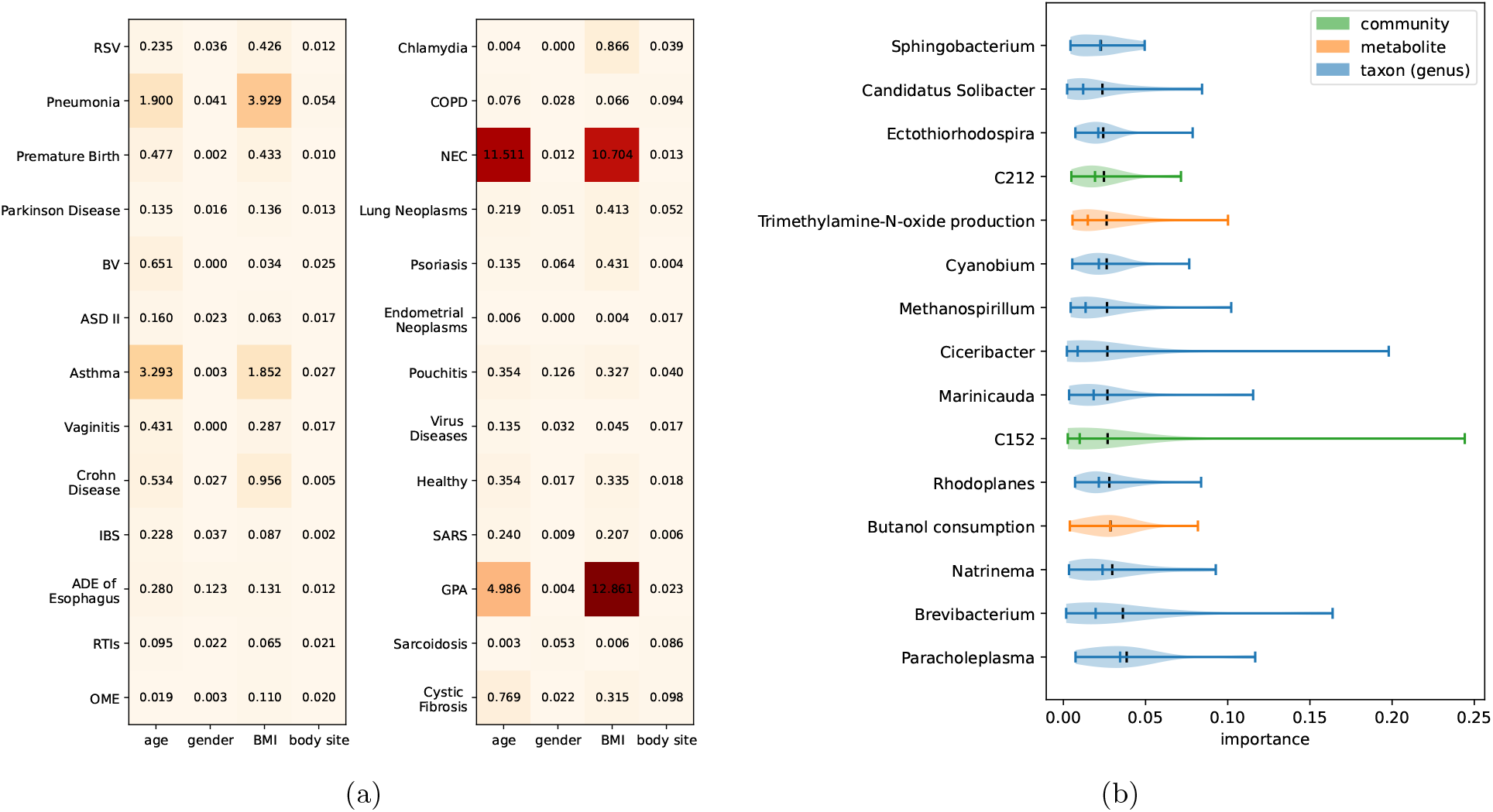
Example interpretations of the predictive models. (a) Impacts of metadata on disease prediction. (b) Impacts of the meaningful hidden nodes on the age prediction.

Figure 5a shows the impacts of metadata on disease prediction. Most of the impact values are non-zero, suggesting that the metadata consistently enhances prediction accuracy across all diseases. Among the metadata we used, age and BMI have large impacts on the prediction of a few diseases including pneumonia, asthma, necrotizing enterocolitis (NEC), and granulomatosis with polyangiitis (GPA). We note that one needs to be cautious with the interpretation for GPA, as GPA was one of the diseases that were not well predicted by MicroKPNN-MT (see Figure 3). For the two respiratory diseases, the metadata has a larger impact on the prediction of asthma than COPD. We note that age and BMI showed similar patterns in their contribution to disease prediction, which could partially be caused by the correlation between age and BMI. Figure 5b shows the impacts of the metabolite, taxon, and community nodes on age prediction. It lists the top 15 contributing hidden nodes (in the Maskedlinear layer that have biological meaning) to the age prediction, including two metabolite nodes, two community nodes, and 11 genera. Interestingly, the genus that is ranked second is *Brevibacterium*, which contains species such as *Brevibacterium epidermidis* that are typically found on the skin. This result is consistent with a previous study, which showed that using skin microbiome resulted in the most accurate prediction of age compared to microbiome from other body sites [26].

## Discussion

We developed MicroKPNN-MT and its application to a large collection of human microbiome datasets in mBodyMap showed that our new method achieved promising results for disease predictions based on microbiome data. In addition, it provided predictions of missing metadata of the microbiome samples, which we anticipate could be utilized by other applications, considering that metadata are largely missing for the existing microbiome datasets. The comprehensive architecture we designed for MicroKPNN-MT enables the model to learn intricate relationships within microbiome samples by combining information from species abundance, prior knowledge about microbial interactions, taxonomic relationships, and community structures. The use of customized layers and metadata-specific encoders contributes to the model’s interpretability and performance across multiple prediction tasks.

Testing on samples from unseen projects showed a substantial reduction in the accuracy of the models due to the heterogeneous nature of the microbiome datasets. The observed divergence in performance between the models using and without using the metadata, especially on the unseen samples, showcased the significance of using metadata to enhance the model’s robustness beyond the confines of the training data. These results further emphasize the potential of metadata-driven approaches to improve predictive outcomes in real-world healthcare applications.

We used microbiome datasets collected in mBodyMap to train and test the models. The mBodyMap dataset is comprehensive containing tens of thousands of samples, and the datasets from different projects were reanalyzed using the same procedures. Limitations include that there are other methods that can be used to infer species profiles from metagenomes, and studies have shown that the choice of bioinformatics analyses could have an impact on the utility of microbiome data [5].

We anticipate that MicroKPNN-NT can be further improved with additional optimization of the models and adding additional functionality. We observed that the model for both disease and BMI prediction actually resulted in slightly worse performance for disease prediction compared to the model that did disease prediction only (see Table 1). One direction of optimization is to try different combinations of the metadata and see how that changes the performance. Second, MicroKPNN-NT can be improved for predicting multiple diseases (a person could have disease A and disease B). Currently, MicroKPNN-MT only predicts one class for the health status prediction (healthy or one of the other 25 diseases). Finally, it is important to further develop MicroKPNN-MT so that it can provide confidence for prediction so if a microbiome dataset is derived from an individual who doesn’t have any of the included phenotypes, MicroKPNN-MT prediction should be able to reflect that.

